# Copy Number Variation at 16p11.2 Imparts Transcriptional Alterations in Neural Development in an hiPSC-derived Model of Corticogenesis

**DOI:** 10.1101/2020.04.22.055731

**Authors:** Julien G. Roth, Kristin L. Muench, Aditya Asokan, Victoria M. Mallett, Hui Gai, Yogendra Verma, Stephen Weber, Carol Charlton, Jonas L. Fowler, Kyle M. Loh, Ricardo E. Dolmetsch, Theo D. Palmer

## Abstract

Microdeletions and microduplications of the 16p11.2 chromosomal locus are associated with syndromic neurodevelopmental disorders and reciprocal physiological conditions such as macro/microcephaly and high/low body mass index. To facilitate cellular and molecular investigations of these phenotypes, 65 clones of human induced pluripotent stem cells (hiPSCs) were generated from 13 individuals with 16p11.2 copy number variations (CNVs). Cortical neural progenitor cells derived from these hiPSCs were profiled using RNA-Seq, which identified alterations in radial glial gene expression that precede morphological abnormalities reported at later neurodevelopmental stages. Moreover, a customizable bioinformatic strategy for the detection of random integration and expression of reprogramming vectors was developed and leveraged towards identifying a subset of “footprint”-free hiPSC clones that are available by request from the Simons Foundation Autism Research Initiative. This publicly available resource of 65 human hiPSC clones can serve as a powerful medium for probing the etiology of developmental disorders associated with 16p11.2 CNVs.

## INTRODUCTION

Copy number variations (CNVs) play a role in the etiology of various neuropsychiatric disorders including intellectual disability ^1^, developmental delay ^1^, congenital malformations ^2^, autism spectrum disorder (ASD) ^3, 4^, schizophrenia (SCZ) ^5, 6^, bipolar disorder (BD) ^7^, and recurrent depression ^8^. Microdeletions and microduplications of a 593kb region of chromosome 16p11.2 have been implicated as a penetrant risk factor in the etiology of neurodevelopmental disorders including ASD and SCZ ^9, 10^. This chromosomal region spans 29.4–32.2 Mb in the reference genome (GRCh37/hg19) and encompasses 29 genes, of which 25 are protein coding ^9, 11, 12^. While estimates of prevalence vary, multiple studies report the presence of a deletion or duplication at the 16p11.2 chromosomal locus in approximately 1% of autistic individuals ^3, 9, 11, 13^ and between 0.01% and 0.1% of the general population ^9, 11, 13^. A meta-analysis of seven studies suggests an overall prevalence of 0.76% 16p11.2 CNVs among idiopathic ASD probands ^14^.

The behavioral and physiological phenotypes associated with the 16p11.2 CNV include reciprocal, shared, and gene dosage-dependent abnormalities [**Figure 1A**]. Both microdeletions and microduplications of 16p11.2 are associated with ASD, although ASD represents a greater proportion of the diagnoses associated with 16p11.2 deletion ^14-16^. Conversely, the risk of SCZ is greater in 16p11.2 microduplication carriers ^5, 10^. Reciprocal neuroanatomical phenotypes of the 16p11.2 CNV include differences in head size and brain volume. Specifically, individuals with a microdeletion of the 16p11.2 locus present with macrocephaly, increased overall gray and white matter volumes, increased cortical surface area, and increased axial diffusivity of white matter tracts such as the anterior corpus callosum and the internal and external capsules ^17-19^. Individuals with a microduplication of 16p11.2 present with microcephaly and corresponding decreases in gray matter, white matter, and cortical surface area ^17, 18^. Recent studies have characterized independent and reciprocal abnormalities in both auditory processing delays and the amplitude of visual evoked potentials among individuals with the 16p11.2 CNVs ^20, 21^. Finally, obesity and hyperphagia are observed among deletion carriers, while low body mass index (BMI) is observed in duplication carriers ^11, 22^.

**Figure 1.**
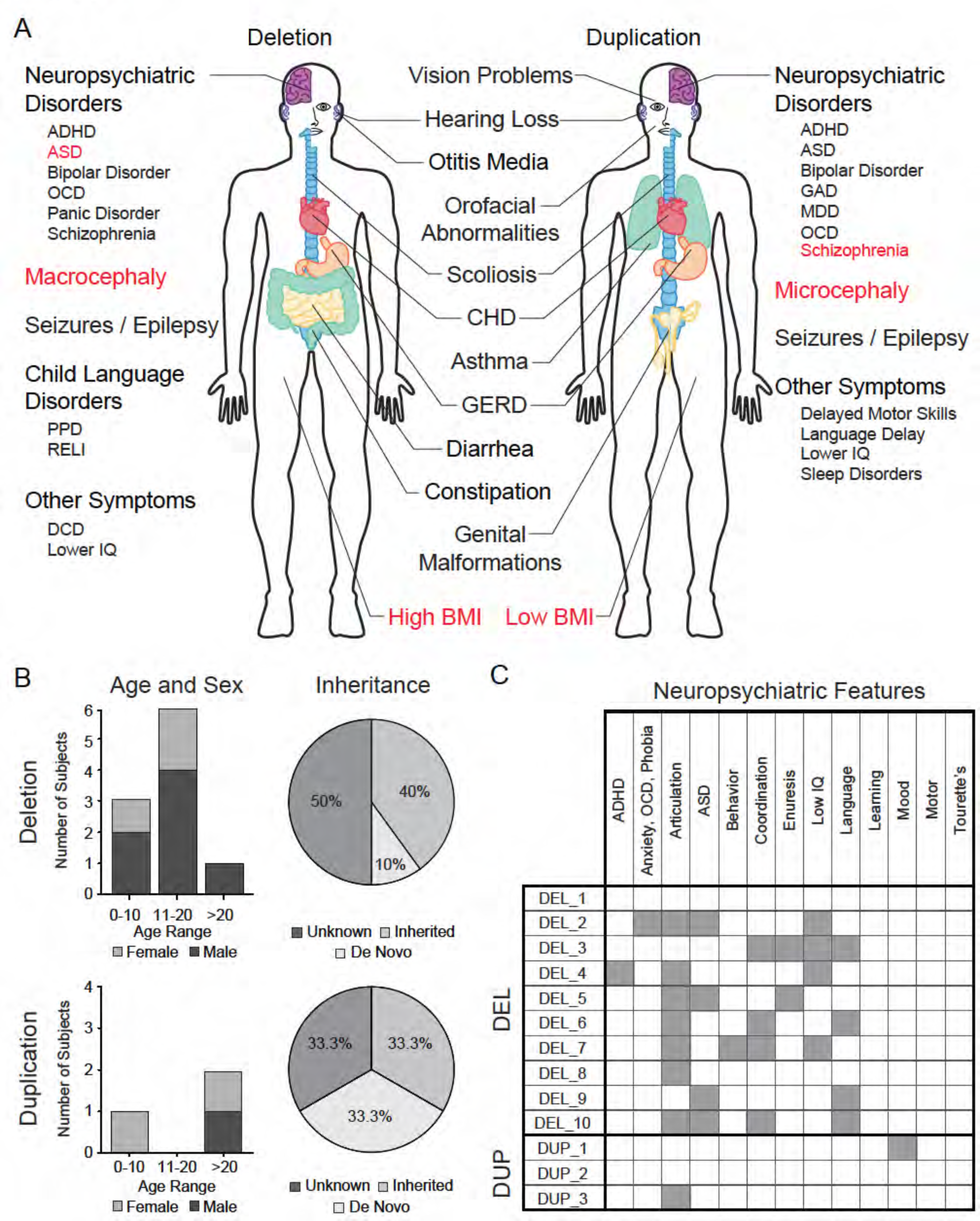
Summary of 16p11.2 CNV clinical features and subject demographics. See also Table S1. (A) Microdeletions and microduplications of the 16p11.2 chromosomal region are implicated in a collection of aberrant behavioral, physiological, and morphological conditions. Common conditions associated with each copy number variant are listed here. Red text indicates reciprocal phenotypes. Abbreviations: ADHD, attention-deficit/hyperactivity disorder; OCD, obsessive-compulsive disorder; PPD, phonological processing disorder; RELI, receptive-expressive language impairment; DCD, developmental coordination disorder; CHD, congenital heart disease; GERD, gastroesophageal reflux disease; GAD, generalized anxiety disorder; MDD, major depressive disorder. (B) A summary of age, sex, and mutation inheritance information for individuals with the 16p11.2 CNV whose fibroblasts were reprogrammed into hiPSCs. (C) Neuropsychiatric attributes in fibroblast donors. Additional neuropsychiatric information exists for each individual (see SFARI VIP database, Table S1). Dark gray boxes indicate positive diagnoses, while white boxes represent negative diagnoses.

The underlying cellular and molecular mechanisms by which neuropsychiatric disorders develop remain largely unknown. Investigators probing the etiology of neurodevelopmental disorders acquired a powerful new tool with the discovery that human somatic cells can be reprogrammed into human induced pluripotent stem cells (hiPSCs) which, in turn, can be differentiated into ectodermal, endodermal, and mesodermal derivatives ^23, 24^. In the years that followed, the field of hiPSC-mediated neurodevelopmental disease modeling, while hampered by intra- and inter-patient variability ^25^, has capitalized on the medium’s unique ability to provide: 1) a model of human development at pathologically-relevant time points, 2) an unlimited source of cells for examination, and 3) tissue-specific cell types which share the genetic background of the somatic cell donor. While initial efforts were focused on monogenic neuropsychiatric disorders ^26-28^, recent work has expanded to include early investigations of gene-environment interactions ^29^ and complex polygenic disorders ^30-34^.

Human iPSCs have offered a novel window into human-specific alterations in neurodevelopment for a number of disorders. Previous work has shown that neurons derived from 16p11.2 patient hiPSCs exhibit abnormal somatic size and dendritic morphology ^35^. Interestingly, transcriptional and phenotypic abnormalities have also been observed in hiPSC-derived cortical neural stem cells derived from individuals with idiopathic ASD ^36, 37^. The 16p11.2 deletion has widespread effects on signaling pathways that underpin critical neural progenitor functions, including proliferation and fate choice ^38^, yet 16p11.2 CNV-related alterations in gene expression patterns have not been examined in early neuroepithelial precursors (radial glia), the stem and progenitor cells of the developing cortex.

Here, we report the derivation, neural differentiation, and transcriptomic characterization of hiPSCs reprogrammed from donors with the 16p11.2 CNV. In total, 65 hiPSC clones were generated from 13 donors. Ten donors harbor a microdeletion at the 16p11.2 chromosomal locus and 3 donors carry a microduplication. All lines reported here are available through the Simons Foundation Autism Research Initiative (SFARI) (https://sfari.org/resources/autism-models/ips-cells). As a cautionary note, we also report on the relatively high frequency at which hiPSC clones were found to contain randomly integrated reprogramming vectors, in spite of the use of non-integrating episomal reprogramming strategies. In many clones, this led to the inappropriate expression of reprogramming genes in neural progenitors, which had a more penetrant impact on genome-wide gene expression patterns than the 16p11.2 CNVs. Moreover, we report that only a subset of genes within the 16p11.2 CNV locus are expressed in early neural progenitor cells, and 14 of these genes show significant reduction in RNA abundance in clones that carry the 16p11.2 deletion. We also show that these alterations are accompanied by changes in the expression of 93 additional genes that are not located within the 16p11.2 deletion interval, including genes with known relevance in neurodevelopment. These genes impinge on signaling pathways relevant to recently described phenotypic abnormalities in neurons *in vitro* ^35^ and in human carriers *in vivo* ^17, 18^.

## RESULTS

### The 16p11.2 microdeletion and microduplication fibroblast donor population is demographically and phenotypically diverse

hiPSCs were derived from skin fibroblasts isolated from individuals recruited to participate in the Simons Variation in Phenotype (VIP) project. A full spectrum of physiological and neuropsychological evaluations was performed and catalogued throughout their involvement with the project and is available to investigators through SFARI. Here, we present a summary of demographic and diagnostic information for 13 individuals who carry a 16p11.2 CNV and for which at least two hiPSC clones are generated [**Figure 1B, C**].

Within the cohort of ten deletion donors, there are seven males and three females ranging in age from 6 to 41 years (mean 13.8 ± 9.98 yrs) [**Figure 1B**]. Seven of the ten donors are probands, one is the father of a proband, and two are siblings of a proband. It is known that four donors carry *de novo* mutations, and one carries an inherited mutation. Of the three duplication donors, two are female and one is male with ages ranging from 5 to 38 years (23.33 ± 16.80 yrs). Among the three duplication donors, one is a proband and the other two are a mother (of the aforementioned proband) and a father of a proband from whom hiPSCs were not derived. Of the three donors, one carries a *de novo* mutation and one carries an inherited mutation.

A battery of cognitive and psychiatric evaluations was performed on each donor. Diagnoses were made in accordance with criteria established in the fourth edition of the Diagnostic and Statistical Manual of Mental Disorders (DSM-IV). Among the 16p11.2 deletion donors, the most common diagnosis was articulation disorder (seven of ten individuals), followed by ASD (four of ten), an IQ below 70 (four of ten), language disorder (four of ten), and coordination disorder (four of ten) [**Figure 1C**]. None of the 16p11.2 duplication donors were diagnosed with ASD or SCZ, although one individual has articulation disorder while another has mood disorder [**Figure 1C**]. Additional metadata for each donor are included in **Supplementary Table 1**.

### 16p11.2 CNV skin fibroblasts were reprogrammed into hiPSCs

Skin fibroblasts from each 16p11.2 CNV donor were transformed with episomal reprogramming vectors that express SOX2, OCT3/4, KLF4, LIN28, L-MYC, and P53-shRNA ^39^ [**Figure 2A**]. Candidate hiPSC clones were identified by the emergence of tightly packed colonies of cells with high nucleus to cytoplasm ratios and sharp colony margins [**Supplementary Figure 1A, B**]. Immunocytochemical (ICC) staining for four pluripotency markers Nanog, Oct3/4 (also known as Pou5f1), TRA-1-60, and TRA-2-49 confirmed that cells in each clone were >95% positive for each marker [**Supplementary Figure 1C-F**]. hiPSC pluripotency was further verified by directed differentiation of the clones into endoderm, mesoderm, and ectoderm lineages, and by assessing changes in pluripotency and lineage marker expression specific to the three germ layers [**Figure 2B, Supplementary Figure 1G**]. Additional quality control data for each hiPSC clone are included in **Supplemental Table 2**.

**Figure 2.**
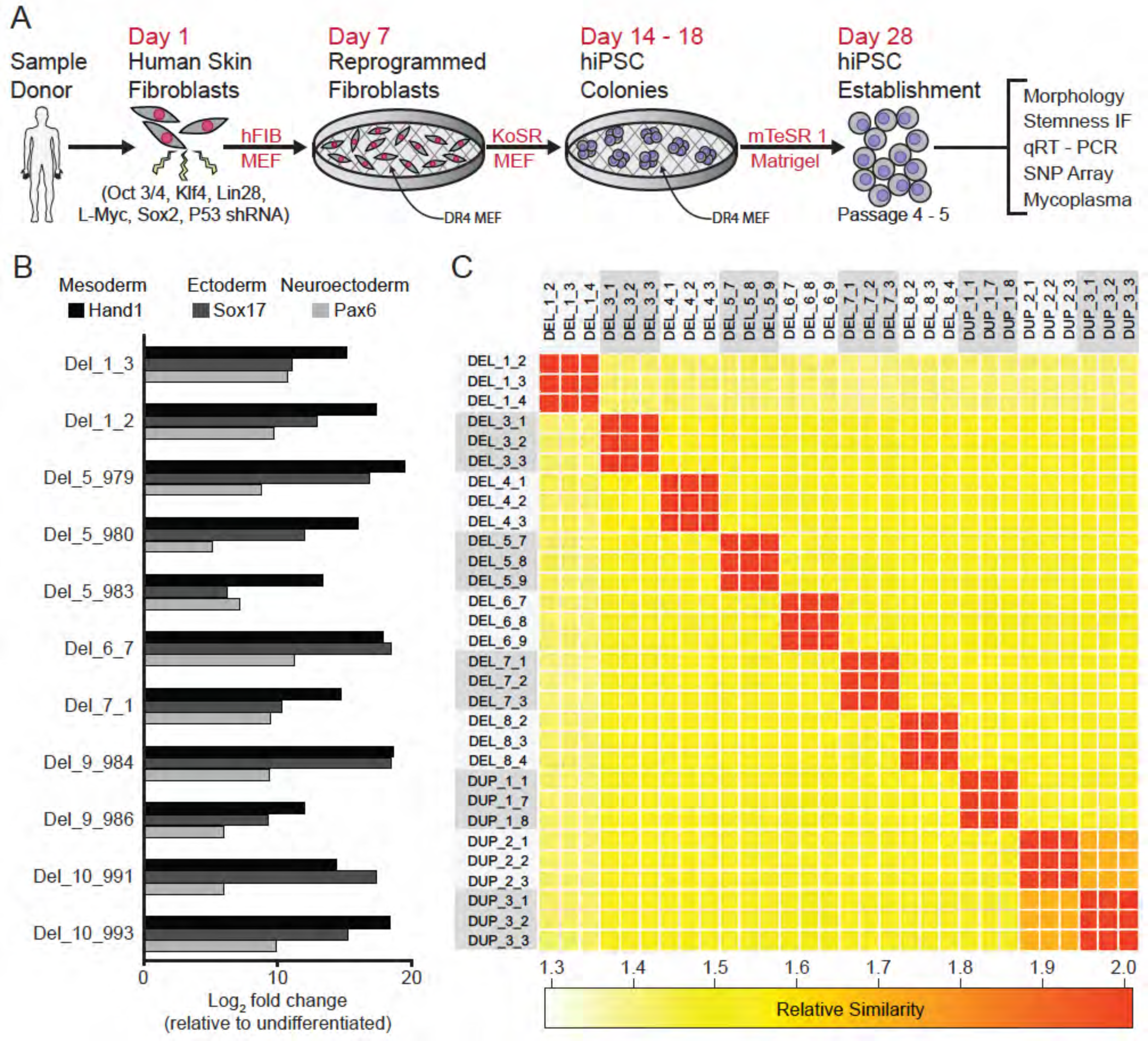
Derivation and validation of 16p11.2 CNV hiPSC. See also Table S1, S2, and S3; Figure S1 and S2. (A) Schematic of episomal reprogramming of human fibroblasts into hiPSCs. (B) qPCR analysis of definitive endoderm (Sox17), lateral mesoderm (Hand1) and neuroectoderm (Pax6) marker expression following directed differentiation. (C) SNP-based similarity matrix illustrating the degree of familial relatedness across a subset of hiPSC clones. Increased similarity between clones is indicated in red. Family members share a larger number of SNPs (orange) than unrelated individuals (yellow).

The majority of hiPSC clones also underwent analysis by SNP array ^40, 41^ to characterize the degree of relatedness between clones, ploidy, and the presence of additional CNVs. All clones were confirmed to be diploid, and, with the exception of two sets of clones from related donors, all clones were unrelated to each other, verifying that archived clones are unique to the respective donor and that inadvertent cross-contamination of cells from independent donors did not occur [**Figure 2C**]. Genetic similarity was confirmed for the two familial donors (DUP_2 and DUP_3) who were known to be mother and daughter. SNP analysis also confirmed that the SNPs proximal to the 16p11.2 CNV breakpoints were consistent with previously reported values, ranging from genomic coordinates 29.4 Mb to 32.2 Mb ^9, 11, 12^ [**Supplementary Table 1**]. Finally, the SNP analysis revealed that CNVs outside of the 16p11.2 locus exist in several patients, which are summarized in [**Supplementary Table 3**].

### 16p11.2 CNV hiPSCs form patterned cortical neural rosettes

40 of the 65 clones were subjected to an adaptation of the monolayer dual-SMAD inhibition protocol^42^ that promotes the formation of dorsal forebrain patterned neural rosettes and a variety of neuronal and glial subtypes [**Figure 3A**]. Flow cytometry confirmed the rapid extinction of Oct4 expression and induction of the radial-glial marker paired box 6 (Pax6) within 7 days [**Supplementary Figure 2A**]. Further differentiation of the cells resulted in the formation of neural rosettes composed of radially arranged radial glia, which is considered to be an *in vitro* recapitulation of both the cellular identity and morphology of radial glia in the developing neural tube ^43, 44^ [**Figure 3B**]. There were no subjective differences between WT and 16p11.2 CNV clones in their ability to form rosettes [**Supplementary Figure 2B**]. After 26 days of differentiation, cells were fixed and stained for a panel of nuclear and cytoplasmic markers to evaluate radial glial cell identities. In time-matched wild-type (WT) control rosettes and 16p11.2 CNV rosettes, Pax6-positive radial glia were similarly arranged in radial clusters [**Figure 3B, Supplementary Figure 2B**]. The establishment of normal epithelial polarity was inferred by the strong apical localization of N-Cadherin (NCad)-positive adherens junctions [**Figure 3B, Supplementary Figure 2B**]. The apical end feet of the radial glia were positive for the tight junction marker zonula occludens-1 (ZO-1) as well as the PAR complex protein atypical protein kinase C zeta (aPKCζ) which is consistent with the establishment of radial glial apical-basal polarity [**Figure 3B, Supplementary Figure 2B**]. Additionally, the mitotic cells, indicated by phospho-histone H3 (pHH3), were found within rosettes proximal to Pericentrin labeled centrosomes which were predominantly located in the center of the rosette, a localization that mimics the apical localization of mitotic cells observed in radial glia *in vivo* ^45^ [**Figure 3B, Supplementary Figure 2B**]. A subset of clones was further differentiated to assess their ability to generate immature neurons. After 45 days of monolayer differentiation, cells exhibited characteristically long, branching neuron-specific class III β-tubulin (Tuj1) positive projections, and a small portion of young neurons were positive for neuronal nuclear protein (NeuN) [**Supplementary Figure 2C**].

**Figure 3.**
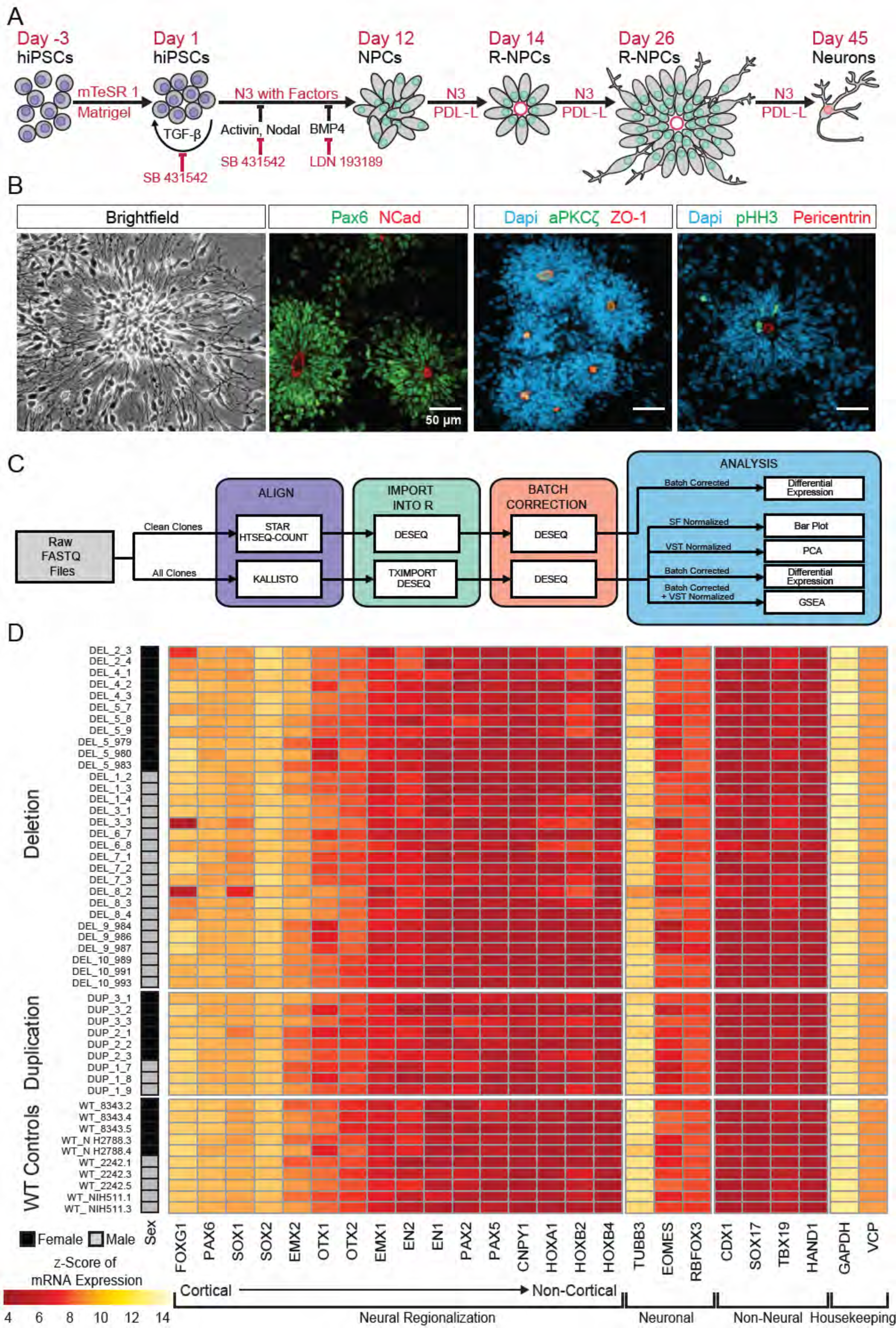
hiPSCs differentiate into cortical neural lineages. See also Table S2 and Figure S3. (A) Schematic of neural differentiation of hiPSCs into cortical progenitor cells and neurons utilizing dual SMAD inhibition. (B) Day 26 neural rosettes show the typical radially arrayed clusters of neural progenitor cells in brightfield micrographs. Rosettes are composed of Pax6-positive radial glia encircling a NCad-positive, ZO-1-positive, and aPKCζ-positive apical adherens complex. Cells currently undergoing M-phase of mitosis, indicated by pHH3, are predominately localized around Pericentrin positive centrosomes at the apical end foot of radial glia (scale bars, 50 µm). Representative rosettes are shown from left to right for WT_8343.2, WT_8343.2, WT_8343.4, and WT_2242.5). (C) Schematic of two bioinformatics pipelines used to characterize hiPSC-derived rosette transcriptomes and validate differentiated cell regional identity. According to the needs of the application, data were normalized by size factor (SF) and by variance stabilizing transformation (VST). A fast and readily replicable pseudo alignment-based pipeline utilizing KALLISTO was later used for detecting and exploring the impact of reprogramming factor integration and expression on the genomic transcriptome. (D) Normalized transcript expression levels of neural regionalization candidate genes generated from mRNA- Seq data, ordered from rostral to caudal cell fates, followed by general neuronal and non-neural cell fates, and housekeeping genes. Sex and Genotype status are indicated on the left, and a visual key for heat map expression values is below.

To more thoroughly evaluate the patterning and differentiation of cells within the neural rosettes, total mRNA was collected from hiPSC-derived neural rosettes after 22 days of differentiation and genome wide transcriptomes were evaluated [**Figure 3C]**. Our initial analysis prospectively examined markers that should be enriched or depleted in neurectoderm of the dorsal telencephalon ^46^. All clones showed strong enrichment for anterior- and dorsal-specific markers, confirming the establishment of anterior cortical identities in culture [**Figure 3D**].

### Reprogramming vector integration induces transcriptional abnormalities

We used principle components analysis (PCA) to evaluate transcriptional variance across the differentiated clones. Surprisingly, PCA showed strong segregation of the clones into two distinct populations that were entirely unrelated to 16p11.2 genotype [**Figure 4A**]. With further analysis, we found that principle components 1 and 2 were associated with elevated expression of Oct3/4, which was subsequently confirmed by quantitative polymerase chain reaction (qPCR) and quantitative real time PCR (qRT-PCR) to be due to the unanticipated random integration, and inappropriate expression, of episomal reprogramming vectors [**Supplementary Figure 3**].

**Figure 4.**
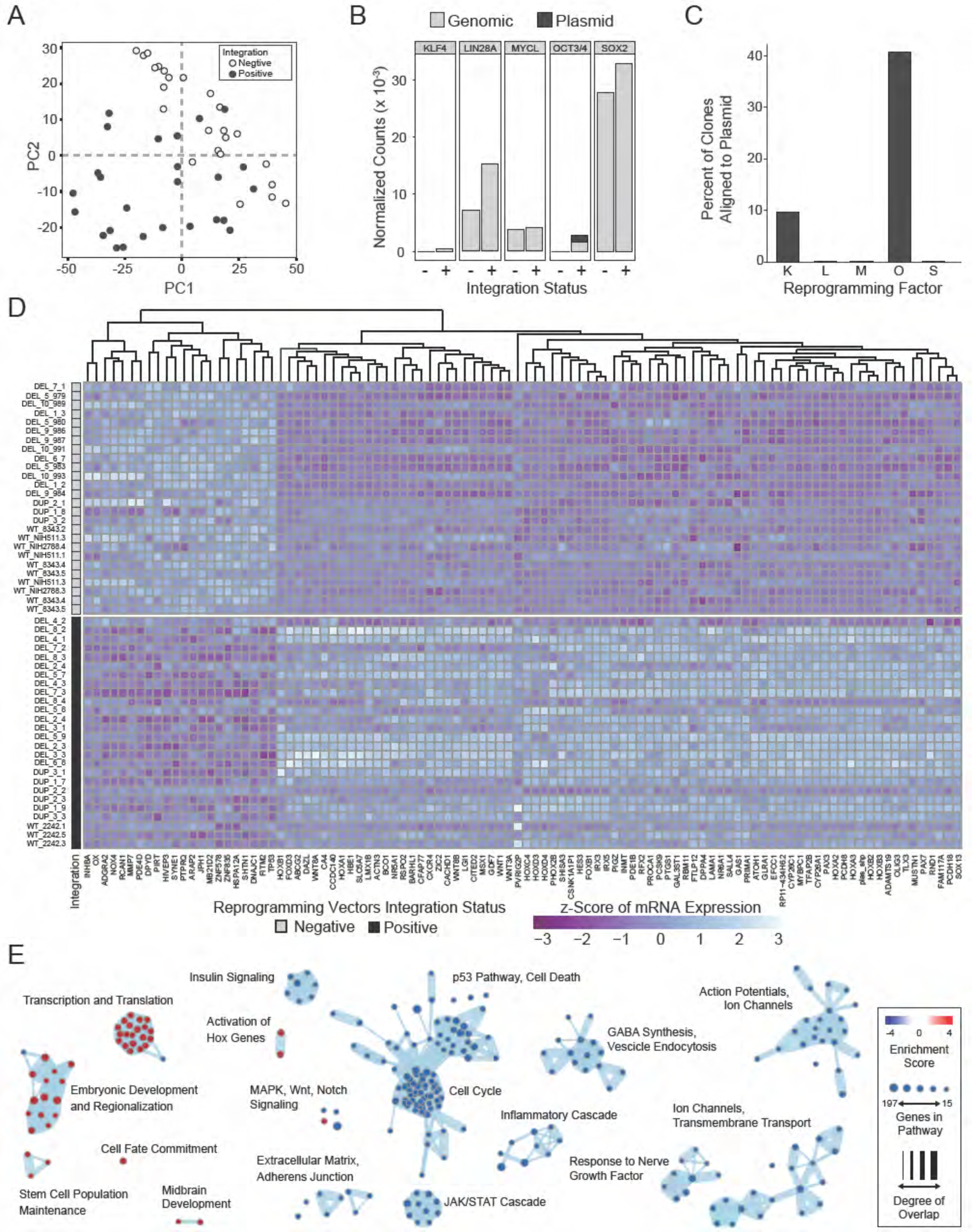
Integration and expression of reprogramming vectors generates pronounced artifacts in the transcriptome. See also Table S2 and S4; Figure S3 and S4. (A) PCA of variance-stabilized count data before batch correction reveals that samples cluster by integration status within the first two PCs. Axes represent the first two principal components (PC1, PC2). (B) Reprogramming factor expression from reads pseudo aligned to the human genome or to plasmid sequences in Int- and Int+ clones. Y-axis represents estimated counts normalized by size factor. The absence of plasmid-aligned transcripts for most genes is indicated by the absence of dark gray segments for each bar (with the exception of Oct3/4). (C) Percentage of total KLF4 (K), LIN28A (L), MYCL (M), OCT3/4(O), or SOX2 (S) counts pseudo-aligned to plasmid in Int+ clones. Y-axis represents the percentage of counts reported in Figure 4B. (D) Heatmap of gene expression represented as Z-scores for the top 100 differentially expressed genes in Int- and Int+ clones as identified with DESeq2. Counts were normalized and scaled using a variance-stabilizing transformation implemented by DESeq, with batch effect correction using limma. Integration status is visualized on the left (integration free clones on top, light gray indicator). (E) GSEA analysis of DESeq output to visualize biological functions potentially impacted by cryptic reprogramming vector integration. Individual nodes represent gene lists united by a functional annotation; node size corresponds to the number of genes in pathway, and color reflects whether the pathway is upregulated (red) or downregulated (blue). Only nodes with significant enrichment in our DESeq output are displayed. The number of genes shared between nodes are indicated by the thickness of their connecting lines. For ease of visualization, individual node labels have been replaced with summary labels for each cluster.

With a novel pseudo-alignment pipeline, we were able to deconvolve counts for a given reprogramming gene to those generated from the genome and those generated from an integrated plasmid [**Figure 4B, C**]. To identify the relative contribution of transcripts from the genome compared to the plasmid integrant, we re- aligned the raw sequencing reads to a composite reference genome consisting of a human reference genome (Ensembl GRCh38.93) and plasmid sequences inserted as extra chromosomes [**Figure 4B**]. With this method, we were able to identify whether the integrated plasmids were transcriptionally active, as well as pinpoint potential transcriptional effects of the integration. Notably, the integration-free hiPSC-derived neural progenitor cells (NPCs) express Lin28a, MycL, and Sox2. Although several clones had integrated plasmids that encode each transcript, the reprogramming vectors did not significantly contribute to the abundance of the respective mRNAs [**Figure 4C**]. Conversely, approximately 10% of KLF4 transcripts detected were expressed from the plasmid and 40% of POU5F1 (OCT3/4) transcripts were plasmid-derived. Although all tested hiPSC clones were competent to differentiate into cortical neural rosettes, we hypothesized that the presence of POU5F1 integrant may affect the self-renewal capacity of the differentiated NPCs ^47-49^. We observed a significant increase (p = 0.03174) in the proportion of pHH3-positive cells in the DEL Int+ differentiated NPCs affected by random integration of the POU5F1 plasmid [**Supplemental Figure 4A**].

To more clearly define the impact of unanticipated episomal vector integration and to determine if clones harboring integrated reprogramming vectors could be used for further analysis, a full RNA-seq analysis was performed on all clones. Differential expression analysis was used to identify alterations due to the integrations. After correcting for the effects of sex, 16p11.2 genotype, and sequencing batch [**Supplemental Figure 4B – E**], we identified 3,612 differentially expressed (DE) genes with adjusted p-value < 0.05, of which 1,739 genes (48.15%) were downregulated in integration-containing (Int+) clones relative to integration-free (Int-) clones. Transcripts attributed via alignment to integrated plasmids were detected and significantly elevated for all three reprogramming plasmids, as well as in transcripts associated with genomic POU5F1, KLF4, SOX2, and LIN28A. There was no statistically significant difference in MYCL. All of these DE transcripts were increased in the INT+ clones, except for the two non-POU5F1 bearing plasmids, which were detected as significantly downregulated in INT+ clones. Given the low baseline expression of these two plasmids relative to the POU5F1-bearing plasmid or the genome-associated transcripts, this DE signature may be dominated by noise generated during the alignment procedure.

The full list of DE genes impinges on major cell functions, such as numerous HOX genes (e.g. HOXA2, HOXB2), regulators of cell morphology (PCDH8, ACTN3), cell cycle (CDKN1A, AKT3, STAT3), and major signaling pathway effectors (e.g. TP53, WNT1). Within the top 100 DE genes [**Figure 4D**], a clear transcriptional difference emerges that seperates Int+ from Int- clones, regardless of 16p11.2 CNV. Although less acute, differences in cell fate acquisition are also observed when clones are sub-divided into Int+ and Int- **[Supplemental Figure 4F**]. To computationally explore the cellular functions that might be implicated erroneously by these transcriptional changes, we applied Genome Set Enrichment Analysis (GSEA) to further probe biologically significant sets of genes that are enriched at the extremes of a ranked list of p-values. At a false detection rate (FDR) < 0.25, over 52 functions were enriched among low p-value/upregulated genes including pathways related to embryonic development and regionalization, HOX gene activation, transcription, and translation [**Figure 4E**]. 355 functions were enriched among low p-value/downregulated genes including pathways related to cell death, cell adhesion, and the JAK/STAT cascade [**Supplementary Table 4**]. Based on these results, we concluded that the transcriptional effects of integration would confound experimentally relevant phenotypes caused by 16p11.2 CNVs, and the Int+ clones were excluded from subsequent analyses. Importantly, an insufficient number of Int- 16p11.2 duplication clones remained (n = 3) for meaningful analysis, and further studies focused exclusively on 16p11.2 deletion clones.

### The 16p11.2 microdeletion affects the transcriptome of hiPSC-derived cortical neural rosettes

RNA-seq data from Int- 16p11.2 deletion (DEL, n = 13) and wild type (WT, n = 7) were re-normalized and analyzed for the potential effects of 16p11.2 deletion in cortical neural rosettes. The known deletion intervals for each of the DEL clones are illustrated in **Figure 5A**. Surrogate variable analysis (SVA) was used to correct for the potential effects of donor sex, clone preparation date, and other batch effects. PCA revealed that the DEL and WT populations clustered according to genotype within the first two principal components after accounting for batch effect correction [**Figure 5B**]. There are 56 canonical gene symbols located in the 16p11.2 locus between positions 28,800,000 and 30,400,000 [**Figure 5C**]. Of these, 14 of the 16p11.2 region genes were differentially expressed, and all were downregulated which is consistent with copy number-dependent effects on mRNA abundance [**Figure 5D**].

**Figure 5.**
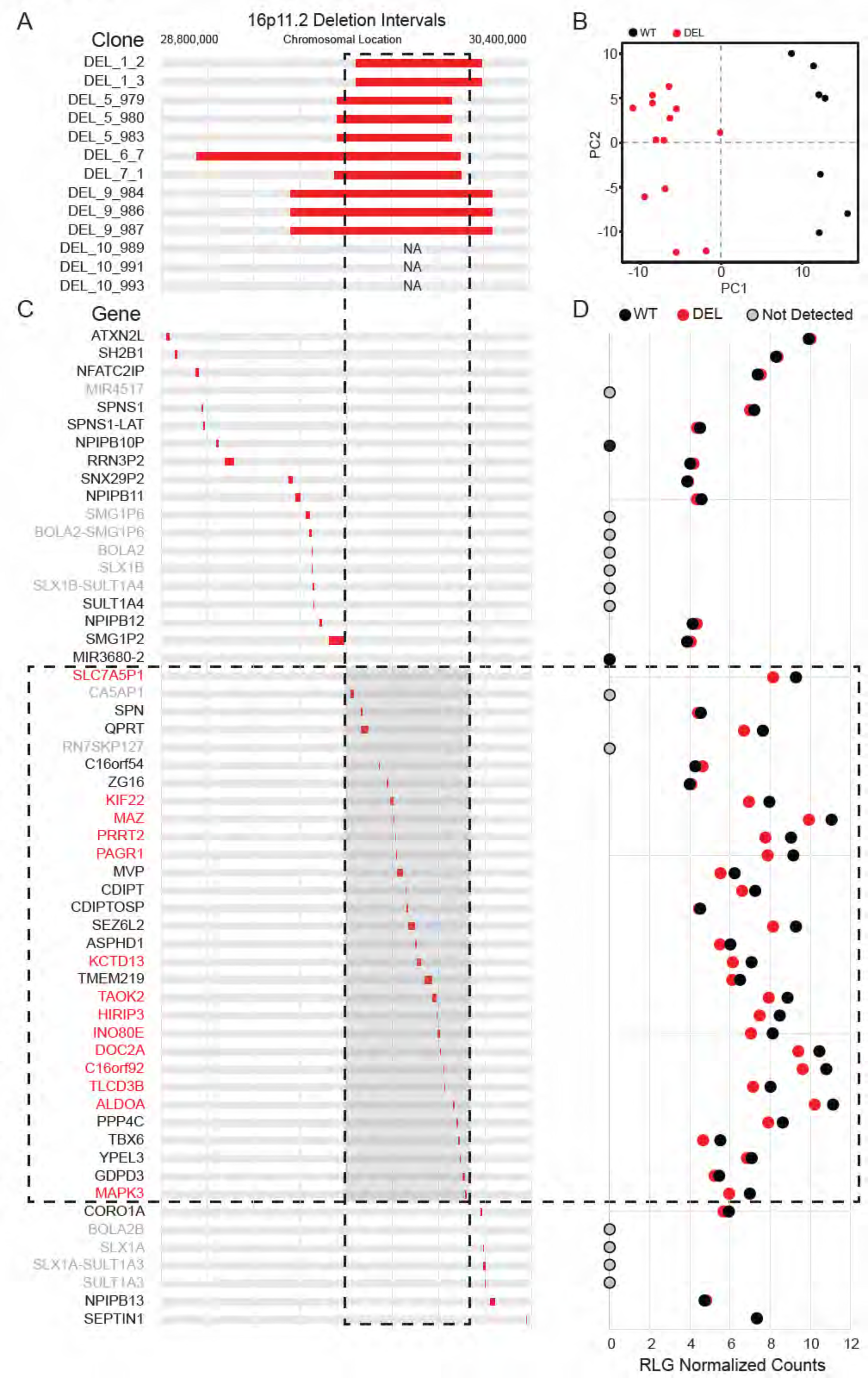
Deletion intervals and differential expression of genes at the 16p11.2 locus. (A) Known deletion intervals in integration-free hiPSC clones that were included in the RNA-seq analysis of differentially expressed genes within the 16p11.2 locus. NA, not available. (B) PCA of variance-stabilized count data after normalization and batch correction reveals that samples cluster by 16p11.2 deletion status within the first two PCs. Axes represent the first two principal components (PC1, PC2). (C) Canonical gene symbols located between chromosome 16 location 28,800,000 and 30,400,000. Transcripts that reach significance as differentially expressed between WT and DEL clones (FDR < 0.05) are indicated in red. Labels for transcripts that were below detection limits are marked in light gray. (D) RLG normalized counts of each transcript within the 16p11.2 interval. WT = black symbols, DEL = red symbols, transcripts not detected = gray symbols.

In total, 107 DE genes were identified in the DEL samples relative to WT (adjusted p-value < 0.05). A full annotated characterization of these genes is provided in the supplement [**Supplementary Table 5**]. The most affected 49 DE genes (1.5-fold or greater change) include all 14 DE genes from the 16p11.2 deletion interval [**Figure 6A**]. A scatterplot of normalized relative mRNA abundance for all transcripts in WT and DEL clones shows relatively tight correlations across genotypes with the deletion interval genes showing a more pronounced change relative to DE genes from other loci in the genome [**Figure 6B**]. Using qPCR we confirmed that a subset (KCTD13, TAOK2, MAPK3 and SEZ6L2) of 16p11.2 region genes that were found to be differentially expressed from our transcriptome analysis were indeed downregulated in each of the DEL samples relative to WT [**Supplementary Figure 5**]. In addition, the individual DEL clones showed strong concordance with the direction of change and approximate amplitude of change for each DE gene within each clone [**Supplementary Figure 6**].

**Figure 6.**
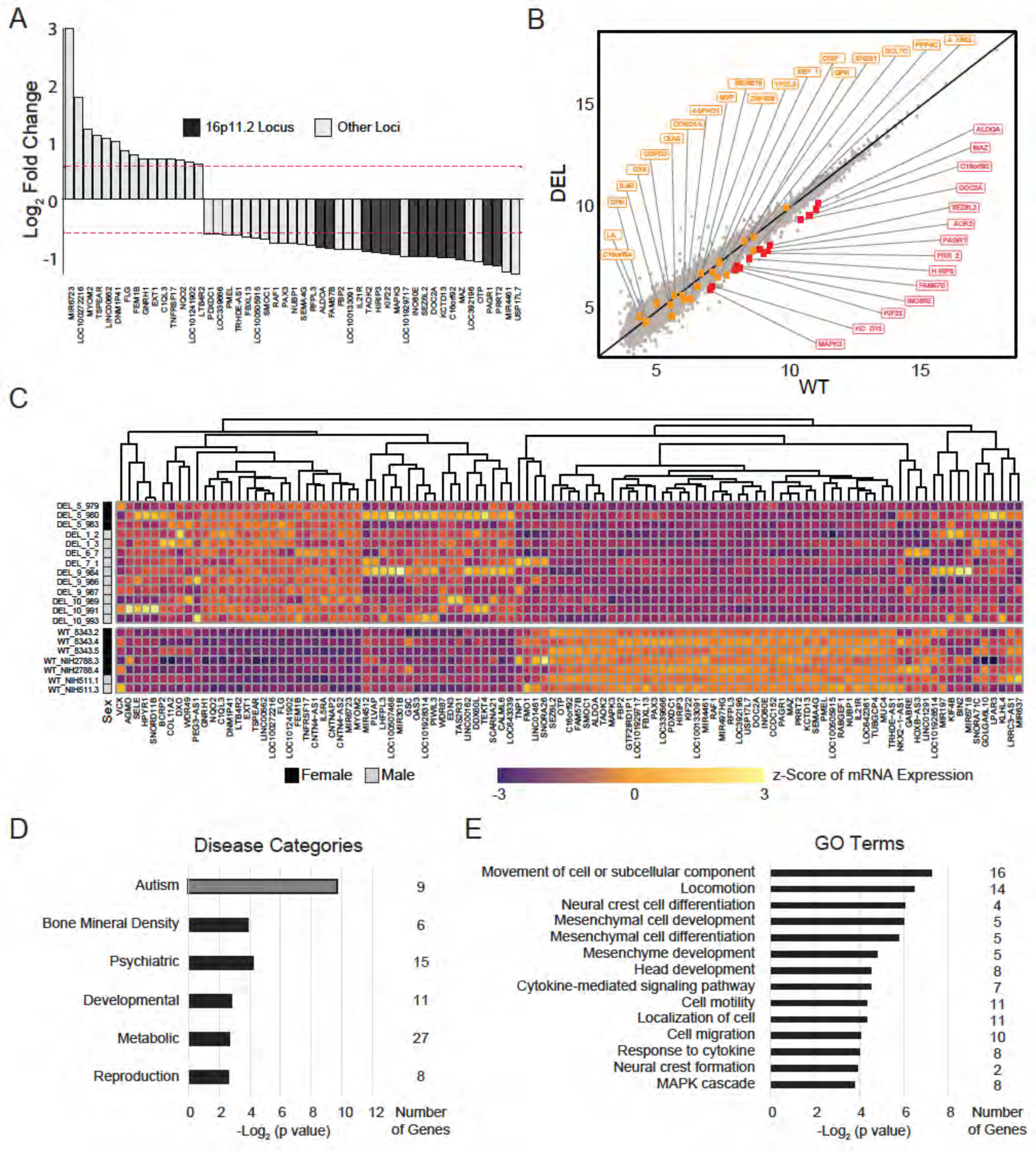
Effects of 16p11.2 deletion on the transcriptome of cortical neural rosettes. See also Table S5 and S6; Figure S5 and S6. (A) Differentially expressed genes that are up- or downregulated at least 1.5-fold. Red lines represent threshold of 1.5-fold change. Genes falling within the 16p11.2 deletion region are highlighted. (B) VST-normalized and batch corrected expression for all genes across all WT clones (X-axis) and DEL clones (Y-axis). Highlighted points represent 16p11.2 region genes that were either called differentially expressed (Red) or not differentially expressed in our pipeline (Orange). (C) Heatmap of gene expression for all the differentially expressed genes identified with DESeq2. Fill values represent counts that have been normalized and scaled using a variance-stabilizing transformation implemented by DESeq, and batch effect corrected using limma and SVA. Sex of the subject is indicated on the left. (D) Disease categories observed in the differentially expressed gene list, ordered by unadjusted p-value, and associated number of genes in each category. Categories that are enriched following Bonferroni correction for multiple hypothesis testing correction are colored dark gray. (E) Gene Ontology (GO) term categories in the differentially expressed gene list, ordered by unadjusted p-value, and associated number of genes in each category.

The entire collection of DE genes [**Figure 6C**] are implicated in a variety of functions. DAVID gene enrichment analysis shows that several disease functions related to neuropsychiatric disorders are associated with more than six genes in the differential expression list [**Figure 6D**]. Additionally, the differentially expressed gene list is statistically significantly enriched for genes associated with Autism, including two genes outside the 16p11.2 region (LHFPL3, CNTNAP2) [**Supplementary Table 6**]. Although the enrichment does not reach significance following multiple hypothesis testing comparison, it is worth noting that additional genes are broadly related to psychiatric disorders (GABRE, GNRH1, FMO1, FLG, SELE, NQO2). Genes relevant to neuroepithelial development are also identified, such as the Erk1/2 MAPK signaling pathway (MAPK3, RAF1, SEMA4G), synapse growth and regulation (CNTNAP2, CALML6, C1QL3, GABRE, PRRT2, SEZ6L2, DOC2A), cytoskeletal organization and cell adhesion (SELE, TEKT4, TUBGCP4, KLHL4), cell cycle (PAGR1, PIWIL3), and fate choice (GSC, OTP, PAX3) [**Figure 6E**]. Although the basic mechanisms that mediate neuroectodermal specification and formation of neural rosettes formation was not measurably altered, the differentially expressed genes within these early stem and progenitor cells strongly suggest that 16p11.2 deletion may impact many functions important to neurodevelopment and neuropsychiatric disease.

## DISCUSSION

Current investigations into the etiology of neurodevelopmental disorders have been limited by an inability to recapitulate the developmental trajectory of the human brain under controlled laboratory conditions. hiPSCs offer a unique opportunity to study these early developmental mechanisms. Here we report the generation and banking of publicly available resource of 65 hiPSC lines with CNVs at the 16p11.2 chromosomal locus. Furthermore, we employ this resource to identify transcriptional alterations that accompany deletion of the 16p11.2 locus in hiPSC-derived NPCs. 16p11.2 CNVs remain one of the most commonly identified variants associated with ASD and, in addition to the previously described links with ASD and SCZ, 16p11.2 CNVs are also implicated in intellectual disability, language disorders, ADHD, motor disorders, and epilepsy^17, 50, 51^.

The hiPSC lines examined here were generated from biospecimens in the Simons Foundation Variation in Phenotype (VIP) Collection and represent lines from 10 deletion donors and 3 duplication donors. These 65 lines are a significant addition to the 14 hiPSC clones recently described by Deshpande *et al*. ^35^ and include donors that are age and sex matched across genotypes. 16p11.2 CNVs are implicated in a spectrum of non-CNS phenotypes, including those of the heart, kidneys, digestive tract, genitals, and bones ^52^. The recruited individuals for this study encompass a broad spectrum of physiological attributes and psychiatric diagnoses. A substantial number of donors is required to verify gene-specific phenotypes in hiPSC models due to intra- and inter-individual variability between clones ^25^. Clone-to-clone variability, in part due to incomplete epigenetic remodeling and the resultant influence of epigenetic memory ^53-55^, may confound conclusions built upon line-to-line comparisons ^56^. Additionally, generating large genomic rearrangements in isogenic control lines through genetic engineering strategies such as the CRISPR/Cas9 system is technically challenging ^57, 58^. Moreover, the availability of a large number of clones provides an opportunity to explore phenotypes that may be relevant to both CNV-specific effects and the influence of an individual’s genetic background on the penetrance of a given phenotype. Importantly, this hiPSC collection contains at least three independently derived clones for most donors. It follows that this resource may significantly improve the field’s ability to identify morphological, physiological, and circuit-level characteristics at different stages of development that contribute to the etiology of 16p11.2-related phenotypes.

The large number of genes within the 16p11.2 region, and their wide-ranging functions, make it challenging to pinpoint the genes that are relevant to neuropsychiatric disease. One of the goals of this work was to determine which of the genes located within the 16p11.2 locus is expressed in early cortical development, as well as which show altered expression following the deletion of one allele. Clones that were free of integrated reprogramming vectors represented genomes from six individuals carrying 16p11.2 deletions. Of these, the deletion intervals were known for five subjects [**Figure 5A**]. We found that all 14 DE genes within the canonical 16p11.2 interval were located within a shorter interval of the chromosome where deletions overlapped for the five subjects with known breakpoints. This clustering to a sub-interval of the locus most likely reflects the fact that the majority of clones analyzed were missing one allele of each DE gene in this region. It is likely that clones with larger deletions may also show copy-number dependent expression for genes outside of the shared interval. For example, it is reasonable to suspect that genes unique to the much larger deletion present in clone DEL_6_7 are downregulated, but that the difference is diluted in the current analysis by clones with normal copy number in this region.

The data presented can be used to confidently identify the genes within the 16p11.2 locus that are expressed in early neural stem and progenitor cells. Of 56 genes within and flanking the deletion intervals of the clones evaluated, 15 of these genes are expressed at or below the limits of detection. Although these genes may play roles at later stages of brain development, they seem unlikely to be relevant for the early patterning and initial establishment of neuronal subtype identity that is being established as NPCs differentiate into neurons.

Conversely, it also stands to reason that the remaining 41 genes are likely to have roles in early cortical development that may be altered when copy number is altered. Of these, 14 are differentially expressed and all are downregulated in DEL: MAZ, DOC2A, PRRT2, PAGR1, TAOK2, KIF22, HIRIP3, ALDOA, C16orf92, INO80E, MAPK3, KCTD13, SEZ6L2, and FAM57B. Their statistically significant downregulation suggests that these 16p11.2 loci are worthy of increased scrutiny as genes that may commonly influence developmental outcomes across individuals that share this deletion interval. Notably, the phenotype may be driven in part by downregulation of 16p11.2 region gene MAPK3, as its protein product Erk1 is known to interact with several other observed DE genes (e.g. SELE, CALML6, RAF1, SEMA4A). This finding supports existing work suggesting that the Erk1 MAPK pathway, which has been shown to modulate the proliferation of cortical progenitors, may drive phenotypic abnormalities associated with the 16p11.2 CNV ^38, 59^. Although we detected a decrease in MAPK3 gene expression in all the 16p11.2 deletion lines [**Supplementary Figure 5A**], we did not observe any changes in NPC proliferation between wild type and deletion lines indicating that MAPK3 may not influence neocortical progenitor cell proliferation in these 16p11.2 patient lines [**Supplementary Figure 4A**]. This is consistent with prior work that also found no significant difference in the proliferation of DEL cells at the NPC stage ^35^.

In addition to the 14 DE genes within the 16p11.2 locus, there are 93 additional genes that are differentially expressed. Fifteen of these genes are identified as being generally related to psychiatric disease, and 9 are more specifically identified as related to autism [**Figure 6D**]. All these genes are listed in the SFARI Gene Scoring Module database ^60^ and are associated with at least five published reports supporting their role as a candidate gene in ASD. Of these genes, CNTNAP2, a neurexin family member that is important excitatory synapse formation and function, is one of the strongest known candidate-genes for ASD risk ^60^. Several additional DE genes are also relevant to excitatory synapses in mature neurons (e.g. PRRT2, CALML6, C1QL3, DOC2A, GABRE, C1QL3, CNTNAP2), and may reflect early perturbations in gene networks relevant to cortical excitatory networks. Previous work does report changes in neurite branching and synapse development in more mature neurons ^35^, and here we find that genes that may be related to these later changes are differentially expressed well before such neurons exist in culture.

Future studies might utilize these early transcriptional features to test whether related functions are affected in mature neurons. For example, the 16p11.2 region gene KCTD13 is important for the maintenance of synaptic transmission in the CA1 region of the hippocampus, with its loss resulting in decreased fEPSP slope and the frequency of mEPSP currents ^61^. The reduction predicted in early progenitor cells might also influence dendrite length, complexity, and spine density. Interestingly, the 16p11.2 region gene TAOK2 has been shown to influence the differentiation, basal dendrite formation, and axonal projection of pyramidal neurons in the cortex with its deletion resulting in abnormal dendrite formation and impaired axonal elongation in mouse brains ^62^. We found the expression of TAOK2 to be reduced across all deletion lines suggesting that reduced TAOK2 and KCTD13 expression in cortical progenitors may impact cortical network development and the maintenance of appropriate synaptic tone in the cortex of autistic individuals.

Other features that the DE gene list may predict include alterations related to movement or locomotion of the cell [**Figure 6E**]. Several DE genes modulate actin and cytoskeletal organization, which could contribute to abnormal cell morphology (e.g. SELE, TEKT4, TUBGCP4, KLHL4). Several of the downregulated genes are related to glucose metabolism and glycogen synthesis (e.g. TLCD3B, FBP2, ALDOA), suggesting that changes in cellular metabolism and energy management may influence outcomes. Finally, we observed that the transcript with the highest fold-change and smallest p-value was aligned the microRNA (miRNA) miR-6723. Although our sequencing protocol was not designed to accurately profile mature miRNAs, the library preparation would capture polyadenylated pri-miRNAs that could later be converted into mature miRNAs^63^. The functions of many of these miRNAs are not well characterized but existing literature supports further investigation. miR-6723 itself is enriched in neurons relative to microglia, astrocytes, and endothelial cells ^64^. Interestingly, it is also downregulated in term placentas of women with chronic Chagas disease, suggesting a role in cellular proliferation or immune response ^65, 66^. Four other miRNAs appear in the differential expression gene list and have roles in proliferation that may impact progenitor function. miR-301B impairs cellular senescence and has a role in the spread of certain cancers ^67, 68^. miR-637 inhibits tumorigenesis, and its downregulation may enhance cell proliferation ^69, 70^.

It is also worth noting that SNP array data show that 4 of the 13 DEL Int- clones contained additional CNVs, including microdeletion in 14q11.1, a microduplication in 14q11.2, and a microdeletion in 7p11.2 in each of the clones. All of these CNVs are potentially associated with neurodevelopmental abnormalities ^71, 72^, and it is possible that these additional CNVs contribute to the transcriptional alterations. However, there is no significant difference in the average fold change of DE gene expression in DEL relative to WT between the subset of clones carrying these additional CNVs and the remaining DEL clones [**Supplementary Figure 5B**]. While it is important to document all background CNVs for future reference, these CNVs are unlikely to account for the transcriptional changes in NPCs reported here.

Finally, we would like to emphasize the importance of screening clones for cryptically integrated reprogramming plasmids, even though published methods emphasize the lack of integration when using “non-integrating” episomal reprogramming vectors. We caution that many of the clones generated in this study and available in the repository contain random integration of reprogramming plasmids that introduce transcriptional artifacts. The impact of cryptic reprogramming vector integration is well illustrated by the clear segregation of Int+ and Int- clones into identifiable clusters within the first two PCs in **Figure 4A**. These transcriptional perturbations could emerge from two possible sources: 1) an increase in reprogramming factor protein product brought about by the insertion of transcriptionally active transcripts, or 2) a disruptive insertion of integrant into a transcriptionally active locus. We note that one Int+ clone (DEL_4_1) has a silenced insertion that contains integrant DNA [**Supplementary Figure 2**] but did not produce detectable transcript that align to the integrated plasmid. Moreover, this Int+ clone has an expression profile that closely resembles those of the Int- clones [**Figure 4D**]. This is consistent with the observed transcriptional effects being primarily related to expression of the reprogramming factors rather than random insertional mutations. The strongest correlation appears to be due to plasmid-based expression of POU5F1 (Oct3/4). This is also suggested in our RT-PCR data, where POU5F transcripts are not detected in clone DE_4_2, which is most similar in DE gene transcriptional patterns observed in the Int- clones [**Supplementary Figure 2**]. The integration and expression of reprogramming vectors is accompanied by the differential expression of more than 3000 genes. Not surprisingly, the integration-associated DE gene list contains 136 genes previously identified as POU5F1 transcriptional targets, representing approximately 16% of genes cataloged in three putative POU5F1 target gene lists ^73^. Given that such cryptic integrations may also exist in other hiPSC lines that have been reprogrammed using non-integrating episomal systems, we provide a customizable RNA-seq analysis pipeline that can detect expression from cryptic plasmids. It is computationally lightweight and can feasibly be performed on a laptop computer. This method provides users with the means to quickly interrogate their data and detect whether similar integration effects should be considered and addressed.

In summary, the collection of 65 human hiPSC clones from 13 donors with either a microdeletion or microduplication at the 16p11.2 chromosomal locus provides an important resource for studying the early neurodevelopmental alterations that may underlie physiological and behavioral features seen in individuals carrying 16p11.2 CNVs. The data presented herein identify several early transcriptomic network perturbations which may act as priming events for later functional alterations in more mature cell types ^37^. A majority of the available clones have been thoroughly evaluated for reprogramming success, ploidy, SNPs, and the capacity to differentiate down the cortical neural lineage. The clones have also been screened and sorted for cryptic integration of the “non-integrating” reprogramming vectors, and data from the subset of footprint-free lines provide several novel candidates for future studies of 16p11.2 deletions. The complete demographic and diagnostic information for each fibroblast donor, and instructions for obtaining the cells themselves, are available through SFARI at https://sfari.org/resources/autism-models/ips-cells. The comprehensive information collected from donors makes this hiPSC resource particularly valuable in understanding potential associations between an *in vitro* phenotype and clinical phenotypes that may be unique to a given individual or shared among individuals with 16p11.2 CNVs.

## MATERIALS and METHODS

### Human hiPSC Generation with Episomal Vectors

The majority of hiPSC available through SFARI were reprogrammed from skin fibroblasts using episomal vectors encoding SOX2, OCT3/4, KLF4, LIN28, L-MYC, and P53-shRNA ^39^. These vectors are pCXLE- hOCT3/4-shp53-F (Addgene, Watertwon, Massachusetts; Plasmid 27077), pCXLE-hSK (Addgene Plasmid 27078), and pCXLE-hUL (Addgene Plasmid 27080). 6 µg of total plasmid DNA (2 µg for each of the three episomal vectors) were electroporated into 5×10^5 fibroblasts with the Amaxaμ Human Dermal Fibroblast Nucleofectorμ Kit (Lonza, Basel, Switzerland; VDP-1001). The fibroblasts were further cultured for 6 days to allow them to recover, and then plated onto mouse DR4 MEF (provided by The Transgenic Mouse Facility at Stanford University). The cells were maintained for 24 hr in growth medium consisting of DMEM/F12 (Thermo Fisher Scientific, Waltham, Massachusetts; 11320033), Fetal Bovine Serum (10%, Thermo Fisher Scientific, 16000044), MEM Non-Essential Amino Acids (NEAA) (1%, Thermo Fisher Scientific, 11140050), Sodium Pyruvate (1%, Thermo Fisher Scientific, 11360070), GlutaMax (0.5%, Thermo Fisher Scientific, 35050061), PenStrep (100 units/mL, Thermo Fisher Scientific, 15140122), and β-mercaptoethanol (55 µM, Sigma Aldrich, M6250). After 24 hours, the culture medium was switched to standard human embryonic stem cell (hESC) medium containing DMEM/F12 (Thermo Fisher Scientific, 11320033), KnockOut Serum Replacement (20%, Thermo Fisher Scientific, 10828028), GlutaMax (0.5%, Thermo Fisher Scientific, 35050061), MEM NEAA (1%, Thermo Fisher Scientific, 11140050), Penstrep (100 units/mL, Thermo Fisher Scientific, 15140122), ß- mercaptoethanol (55 µM, Sigma Aldrich, St. Louis, Missouri; M6250), and human recombinant FGF2 (10 ng/ml, R&D Systems, Minneapolis, Minnesota; 233-FB). After two to three weeks, emerging hiPSC colonies were manually selected based on morphological standards and expanded clonally into hESC-qualified Matrigel Matrix (0.10 mg/mL, Corning, Corning, New York; 354277) coated plates in mTeSR 1 medium (Stem Cell Technologies, Vancouver, Canada; 5851). A subset of the available clones was reprogrammed using the Epi5 Reprogramming Kit (Thermo Fisher Scientific, A15960) according to the kit’s instructions. The hiPSCs were routinely passaged with Accutase (Stem Cell Technologies, 07920) when they reached high confluency and re-plated in mTeSR 1 with RHO/ROCK Pathway Inhibitor Y27632 (Stem Cell Technologies, 72302). Patient donors are identified with a code with one numeral (DEL_1) whereas different lines from the same patient are identified with an extended code (e.g. DEL_1_2, DEL_1_3).

### Expression of Pluripotency Markers in hiPSCs

All cells maintained in monolayer cultures were fixed with 4% paraformaldehyde for 10 min at room temperature. The pluripotency of the derived hiPSCs was tested by immunocytochemical staining for OCT4 (Abcam, Cambridge, United Kingdom; ab27985, RRID: AB_776898), NANOG (R&D Systems, AF1997, RRID: AB_355097), TRA-1-60 (Abcam, ab16288, RRID: AB_778563) and TRA-2-49 (Developmental Studies Hybridoma Bank, TRA-2-49/6E, RRID: AB_528071). The secondary antibodies were goat anti-rabbit IgG conjugated with Alexa-488 (Invitrogen, Carlsbad, California; A-11034, RRID: AB_2576217) and goat anti-mouse IgG conjugated with Alexa-488 (Invitrogen, A-11011, RRID: AB_2534069).

### Episomal Vector Integration and Transcripts Detection

Genomic DNA was extracted from cell pellets using the GeneJET Genomic DNA purification KIT (Thermo Fisher Scientific, K0722) according to supplier protocols. We designed primers using SnapGene software (RRID: SCR_015052) and verified them for the absence of non-specific binding using NCBI Primer-BLAST. Primers were designed to amplify products that spanned part of either one of the reprogramming genes OCT4, KLF4, LIN28, and the flanking WPRE region to prevent primer hybridization and amplification of these endogenous genes in the genomic DNA. To assess the sensitivity of the PCR reaction, plasmid DNA (each of the three plasmids separately) was mixed in with H9 genomic DNA to create a dilution curve at concentrations starting at 2000:1 and reducing by factor of ten to a 2:1 copy number of plasmid to genomic DNA. All PCR reactions were carried out using the Platinum Taq DNA Polymerase High Fidelity kit (Thermo Fisher Scientific, 11304011) according to supplier protocols. 1 µl of genomic DNA (approx. 15 ng) was added to the amplification master mix containing 60 mM Tris-(SO4), 18 mM (NH4)2SO4, 50mM MgSO4, 10 mM dNTPs, 5U/ul of Platinum Taq DNA Polymerase, and the forward and reverse primers. The amplification reaction was carried out using a cycling program of 1 min at 95 °C, followed by 35 cycles of 15 s at 95 °C, 30 s at the annealing temperature specific for each primer set, and 1 min at 68 °C in a BioRad C1000 thermal cycler. Total RNA was extracted from cell pellets using the GeneJET RNA purification kit (Thermo Fisher Scientific, K0731) and cDNA synthesis performed was performed using the High Capacity cDNA Reverse Transcription kit (Applied Biosystems, Foster City, California; 4368814) according to supplier protocols. For the cDNA synthesis, 3 µl of RNA was added to the master mix (10x RT buffer, 100 mM dNTPs, Multiscribe Reverse Transcriptase, 10X primers), and the reverse transcription reaction was performed using a cycling program of 10 min at 25 °C, 120 min at 37 °C, and 5 min at 85 °C. To detect the presence of vector transcripts, 1 µl of cDNA from each clone was used in an amplification reaction with the same set of primers that were used for the detection of vector integration in gDNA. Primer sequences for integrated plasmids and transcripts were as follows: pCXLE- hOCT4-shp53-F (fwd, CAGTGTCCTTTCCTCTGGCCCC; rev, ATGAAAGCCATACGGGAAGCAATAGC; 329 bp product), pCXLE-hSK (fwd, AATGCGACCGAGCATTTTCCAGG; rev, TGCGTCAGCAAACACAGTGCACA; 342 bp), pCXLE-hUL (fwd, CAGAGCATCAGCCATATGGTAGCCT; rev, ACAACGGGCCACAACTCCTCAT; 380 bp). H9 hESC DNA was used as negative controls in all amplification reactions.

### Single Nucleotide Polymorphisms and Copy Number Variants Analyses

The Affymetrix Genome-Wide Human SNP Array 6.0 platform was chosen for SNP and CNV analysis. The SNP array assay was performed and analyzed by CapitalBio Corp., Beijing, China. Genetic markers, including more than 906,600 SNPs and more than 946,000 CNV probes, were included for the detection of both known and novel CNVs. Data were analyzed by a copy number polymorphism (CNP) calling algorithm developed by the Broad Institute (Cambridge, Massachusetts, USA). In addition, sample mismatch analysis was performed with genotyping data of the SNP array to confirm that each clone from a given donor was identical and carried the donor’s original genotype ^40, 41^.

### Differentiation of Human hiPSCs to the Neural Lineage

As previously reported ^42^, hiPSCs were differentiated in N3 culture medium which consists of DMEM/F12 (1x, Thermo Fisher Scientific, 11320033), Neurobasal (1x, Thermo Fisher Scientific, 21103049), N-2 Supplement (1%, Thermo Fisher Scientific, 17502048), B-27 Supplement (2%, Thermo Fisher Scientific, 17504044), GlutaMax (1%, Thermo Fisher Scientific, 35050061), MEM NEAA (1%, Thermo Fisher Scientific, 11140050), and human recombinant insulin (2.5 µg/mL, Thermo Fisher Scientific, 12585014). From Day 1 to 11, the N3 media was further supplemented with two factors: SB-431542 (5 µM, Tocris, 1614) and LDN-193189 (100 nM, Stemgent, 04007402). At Day 12, the differentiating cells were dissociated with Cell Dissociation Solution (1x, Sigma-Aldrich, C5914), passaged onto Poly-D-Lysine (50 µg/mL, Sigma-Aldrich, P1024) and Laminin (5 µg/mL, Roche, Basel, Switzerland; 11243217001) coated plates, and cultured in N3 media without factors until Day 22 when they were passaged again. Between Day 1 and Day 22, media changes were performed daily. However, after Day 22, cells were exposed to media changes every other day with N3 media without factors.

### Differentiation of Human hiPSCs to Endoderm and Mesoderm Lineages

hiPSC cells grow in mTeSR were dissociated into single cells using accutase and plated at density of 25 – 50k cells/cm2 on matrigel coated cell culture plates and subsequently differentiated into endoderm and mesoderm lineages ^74, 75^. For definitive endoderm induction, anterior primitive streak was first specified using 100 ng/ml Activin A (R&D systems, 338-AC-050), 3 µm CHIR (Tocris, 4423) and 20 ng/ml FGF2 (R&D Systems, 233-FB- 01M) in CDM2 basal media. After 24 hr, the cells were washed with DMEM/F12 (1x, Thermo Fisher Scientific, 11320033), and definitive endoderm was induced using 100 ng/ml Activin A and 250 nM LDN (Reprocell, Yokohama, Japan; 04-0074) in CDM2 basal media for 24hr. For lateral mesoderm induction, Midprimitive streak was specified using 30 ng/ml Activin, 16 µM CHIR, 20 ng/ml FGF2 and 40 ng/ml BMP (R&D Systems, 314-BP-050) in CDM2 basal media for 24 hr. After 24hr, cells were washed with DMEM/F12 (1x, Thermo Fisher Scientific, 11320033), and lateral mesoderm was induced using 1µM A8301 (R&D Systems, 2939) 30 ng/ml BMP and 1 µM C59 (Tocris, Bristol, United Kingdom; 5148) for 24hr in CDM2 basal media. On the third day, cells were lysed for RNA collection and purification.

### RNA Extraction, Reverse Transcription and qPCR

RNA was collected from adherent wells grown in individual wells of a 24 well cell culture plate using the Qiagen RNeasy kit as per the manufacturer’s instructions with an added intermediate step of on-column DNA digestion to remove genomic DNA. About 10-100 ng of RNA was used for reverse transcription (High capacity cDNA reverse transcription kit, Applied Biosystems, 4368814) as per the kit manufacturer’s instructions. cDNA was then diluted 1:10 and was used for each qPCR reaction in a 384 well format. To assess gene expression of endoderm, mesoderm and ectoderm lineages, in each individual qPCR reaction, 5 µl of (2x) SYBR green master mix (SensiFAST SYBR kit, Bioline, London, United Kingdom; BIO- 94005) was used and combined with 0.4 µl of a combined forward and reverse primer mix (10 uM of forward and reverse primers in the combined master mix) and the reaction was run at Tm of 60 °C for 40 cycles. qPCR analysis was conducted by the ΔΔCt method and the expression of each gene was internally normalized to the expression of a house keeping gene (Actb) for the same cDNA sample. For all differentiated cells, expression of the lateral mesoderm marker (Hand1), definitive endoderm marker (Sox17), neuroectoderm marker (Pax6) and pluripotency marker (Nanog) was compared to the expression of these markers from the same undifferentiated cell line. The primer sequences for the lineage markers are as follows, Actb (fwd: AGAGCTACGAGCTGCCTGAC, rev: AGCACTGTGTTGGCGTAGAC), Hand1 (fwd: GTGCGTCCTTTAATCCTCTTC, rev: GTGAGAGCAAGCGGAAAAG), Sox17 (fwd: CGCACGGAATTTGAACAGTA, rev: GGATCAGGGACCTGTCACAC), Pax6 (fwd: TGGGCAGGTATTACGAGCTG, rev: ACTCCCGCTTATACTGGGCTA). For the validation of the DESeq interval genes, RNA from cells differentiated for 22 days *in vitro* into neuroectoderm and subsequently patterned to a dorsal telencephalic identity was used for reverse transcription and qPCR. Genes with largest log2 fold change and most significant p values from the DESeq analysis (TAOK2, Taqman assay Hs00191170_m1; SEZ6L2, Taqman assay Hs03405581_m1; MAPK3, Taqman assay Hs00385075_m1; KCTD13, Taqman assay Hs00923251_m1) compared to wild-type lines were selected. To assess the expression of the 16p interval genes, in each individual reaction, 5 µl of (2x) Taqman fast advanced master mix was combined with 0.5 µl of the appropriate Taqman gene expression assay probe. qPCR analysis was conducted by the ΔΔCt method and the expression of each gene was internally normalized to the expression of a house keeping gene (Actb) for the same cDNA sample. The expression of the assayed genes was compared to the average expression of 6 wild type hiPSC clones.

### Expression of Neural Rosette and Immature Neuronal Markers in Monolayer Cultures

Day 26 differentiated neuroepithelial cells were evaluated via immunocytochemical staining of Pax6 (Biolegend, San Diego, CA; 901301, RRID: AB_2565003), NCad (BD Biosciences, San Jose, California; 610920, RRID: AB_2077527), ZO-1 (InVitrogen, 33-9100, RRID: AB_2533147), aPKCζ (Santa Cruz, Dallas, Texas; sc-216, RRID: AB_2300359), Pericentrin (Abcam, ab28144, RRID: AB_2160664), and pHH3 (Santa Cruz, sc-12927, RRID: AB_2233069). Day 45 immature neurons were stained for Tuj1 (Abcam, ab78078, RRID: AB_2256751) and NeuN (Abcam, ab177487, RRID: AB_2532109). The secondary antibodies were donkey anti-rabbit IgG conjugated with Alexa-488 (Jackson, Bar Harbor, Maine; 711545152, RRID: AB_2313584), donkey anti-mouse IgG conjugated with Cy3 (Jackson, 715165150, RRID: AB_2340813), and donkey anti-goat IgG conjugated with Cy5 (Jackson, 705175147, RRID: AB_2340415). Quantification of pHH3 was performed using ImageJ (RRID: SCR_003070) and Adobe Photoshop (RRID: SCR_014199), while all statistical evaluations were performed with GraphPad Prism v7.04 (RRID: SCR_002798).

### Flow Cytometry

Cells were dissociated by treatment with 0.5 mM EDTA (Thermo Fisher Scientific, 15575020) in Phosphate-Buffered Saline (PBS) (Thermo Fisher Scientific, 10010023) for 5 minutes, or Cell Dissociation Solution (1x, Sigma-Aldrich, C5914) for 20 minutes, washed with PBS, fixed in 4% paraformaldehyde for 10 minutes, and washed again with PBS. Cells were then permeabilized with 0.1% Triton X-100 (Thermo Fisher Scientific, 85111) and stained with conjugated primary antibodies for 30 minutes at manufacturer-recommended concentrations at 4°C in the dark. Antibodies used were anti-Pax6 Alexa 647 (BD Biosciences 562249), and anti-Oct4 Phycoerythrin (PE) (BD Biosciences 560186). Isotype matched controls with corresponding fluorchrome conjugates were acquired from BD Biosciences. Cells were analyzed using a BD Aria II.

### RNA Sequencing

Day 22 cortical NPCs were lysed using TRIzol (Thermo Fisher Scientific, 15596026) and stored in -80 °C until. The samples were then delivered to the Stanford Functional Genomics Facility (SFGF) where they were multiplexed and sequenced using paired end mRNA sequencing on an Illumina NextSeq 500. Raw data were demultiplexed at the sequencing facility. RIN scores for generated libraries were nearly all above 9.0 (range: 7.8 – 10.0). Reads were 75 basepairs in length, and samples were sequenced with a targeted effective sequencing depth of 40,000,000 reads per sample. Given the high quality of each read, as established by a per-base sequence quality greater than 28 in FastQC (v0.11.6, RRID: SCR_014583), no further read trimming was performed.

### Assessment of the Impact of Reprogramming Factor Integration

The .fastq files produced by RNA-seq were aligned using kallisto (v0.43.1) ^76^ to a unique index composed of a human reference transcriptome (Ensembl GRCh38.93) and plasmid sequences. The mean number of reads processed per sample was 102,963,875 (range: 65,427,345 – 193,569,749), and these pseudoaligned to a reference mRNA at a mean rate of 76.34%. By adding plasmid sequences as additional ‘chromosomes’, we were able to investigate whether sequencing reads aligned to non-genomic as well as genomic DNA. Estimated counts produced by kallisto for each were imported into RStudio for analysis with R using the tximport function (v.1.8.0) ^77^ and imported into a DESeq2 object for further analysis. Genes with one or fewer detected reads across samples were filtered out to expedite computation. Due to the polycistronic nature of the transcripts produced by plasmids, the same expression value for a given plasmid was considered associated with each of its component reprogramming factor genes. PCA plots were produced using the plotPCA function of the DESeq2 (v1.20.0, RRID: SCR_015687) package ^78^, barplots produced with ggplot2 (v3.0.0, RRID: SCR_014601) ^79^, and heatmaps produced with pheatmap (v1.0.10, RRID:SCR_016418) ^79^. Differential gene expression was calculated using DESeq2 and reported p-values represent FDRs calculated using Benjamini-Hochberg multiple hypothesis testing correction provided by that software pipeline. The threshold for gene significance was set at 0.05. Several batch effect variables were included in the design matrix for the analysis (i.e., patient Sex, patient CNV status, and sequencing day) to ensure that this analysis focused on the transcriptional effects of integration and not a potential batch effect. Pheatmap z-scores are calculated by subtracting the mean from each input expression value (centering) and dividing by the standard deviation (scaling). To investigate potential biological mechanisms impacted by reprogramming factor integration, the genes were ranked according to a metric representing statistical significance combined with the direction of fold change. Using this scheme, the genes were effectively ranked such that the first and the last entries in the list represented the smallest p-value upregulated and smallest p-value downregulated genes, respectively. The resulting list was submitted to GSEA (RRID: SCR_003199) to identify functional gene modules enriched at the top or the bottom of the list with FDR < 0.25. A subset of the resulting enriched modules was visualized using the EnrichmentMap (RRID: SCR_016052) plugin for Cytoscape (RRID: SCR_003032) ^80^. A list of 16p11.2 region genes was acquired by exporting all annotated transcripts in the region from the USCS Genome Browser (RRID: SCR_005780).

### Differential Expression Analysis

A subset of the .fastq files from the previous analysis corresponding to Int- clones were re-aligned using STAR (v2.5.3a, RRID:SCR_015899). The aligned reads were counted using htseq-count. The mean number of uniquely mapped reads for each sample when imported into R was 83,387,539 reads per sample (range: 54,003,338 - 164,989,807). Given that this was a paired-end sequencing experiment, the effective read depth was a mean of 41,693,770 reads per sample. These raw data were batch corrected using SVA (v3.28.0) ^81^, and analyzed using DESeq2 for differential expression analysis.DESeq2 uses the given data to model each gene as a negative binomial generalized linear model, and tests for differential expression using the Wald Test and Benjamini-Hochberg adjusted p-values. For visualization of batch effect corrected data, data were transformed using LIMMA (v3.36.5) ^82^. Shrunken log2-fold changes are used according to the recommendation in DESeq2 to account for fold change inflation in low expression genes. Heatmaps are generated using the R package pheatmap, which created a dendrogram of gene similarity using kmeans clustering. Expression values from RNA-Seq data aligned using STAR were also used to visualize fate marker expression *in vitro*. Gene ontology enrichment analysis was performed using DAVID (RRID:SCR_001881) ^83, 84^. Gene ontology enrichments were ordered by unadjusted p-values, and the top entries plotted. An indication was added where an enrichment reached statistically significant enrichment following Bonferroni-adjusted p-values.

### Distribution of Materials

The distribution of hiPSCs is by permission from The Simons Foundation Autism Research Initiative. Additional information on the Variation in Phenotype Project and the process for requesting biospecimens can be found at https://sfari.org/resources/autism-models/ips-cells. Code for bioinformatic analyses in this paper are available at www.github.com/kmuench/16presource (DOI: 10.5281/zenodo.1948176).

## SUPPLEMENTARY INFORMATION

Supplementary information is available online through eLife.

## ACKNOWLEDGMENTS

We thank the 16p11.2 CNV donors and their families for their essential contributions to this resource, as well as Sara Walton for her efforts in expanding and cryopreserving of many of the clones received from the SFARI collection. We additionally thank the Stanford Functional Genomics Facility for their assistance in generating the RNA-seq data analyzed in this manuscript. This work was supported by the National Institute of Mental Health (NIMH) grant 1R01MH108660 to T.D.P.

## AUTHOR CONTRIBUTIONS

Conceptualization, J.G.R, K.L.M., H.G., T.D.P; Methodology, J.G.R., K.L.M., H.G., A.A; Validation, J.G.R., H.G., A.A., V.M.M.; Investigation, J.G.R., K.L.M., H.G., A.A., Y.V., V.M.M., S.W., J.L.F., C.C.; Formal Analysis, J.G.R., K.L.M., A.A., V.M.M; Visualization, J.G.R., K.L.M., H.G., A.A., V.M.M.; Writing – Original Draft, J.G.R., K.L.M.; Writing – Review and Editing, J.G.R., K.L.M., A.A., V.M.M., K.M.L., T.D.P.; Supervision, T.D.P.; Funding Acquisition, T.D.P.

## CONFLICT OF INTEREST

Ricardo E. Dolmetsch has received compensation from Novartis as a part of its executive board and may use a subset of these lines in research. The remaining authors declare no competing interests.

## FIGURE TITLES and LEGENDS

## SUPPLEMENTARY INFORMATION TITLES and LEGENDS

**Supplementary Table 1. Demographic, diagnostic, and breakpoint information**.

Abbreviations: NA, not applicable; ND, not determined.

**Supplementary Table 2. Pluripotency, vector silencing, and neural competency information**.

(A) Subset summary of quality control data for hiPSC reprogramming and differentiation.

(B) Entire quality control data for hiPSC reprogramming and differentiation.

Abbreviations: NA, not applicable; ND, not determined. Plasmids used for the PCR targeted a portion of the plasmid-specific WPRE region.

**Supplementary Table 3. CNVs located outside the 16p11.2 chromosomal locus**.

SNP array analysis revealed hiPSC-clone-specific CNVs throughout the genome. Their copy number, location, and size are listed.

**Supplementary Table 4. Gene sets significantly enriched in ranked list of DESeq2 output Comparing Int+ and Int- Clones**.

All genes submitted to DESeq2 were ranked according to the -log10 of adjusted p-value and the sign of their fold change, such that the top of the list represented upregulated and significant genes, and the bottom of the list downregulated and significant genes, with non-significant and high p-value genes towards the middle. Gene sets were generated by GSEA and additional annotation provided by Enrichment Map.

**Supplementary Table 5. Differentially expressed genes in 16p11.2 deletion clones relative to control clones**.

Annotated differentially expressed gene list generated by DESeq2. Genes are ranked by adjusted p-value.

**Supplementary Table 6. DAVID gene ontology enrichment analysis**.

Gene ontology enrichments were ordered by unadjusted p-values.

**Supplementary Figure 1. Fibroblast reprogramming and pluripotency validation**.

(A) Bright field representative image of Day 18 clone of reprogrammed hiPSC surrounded by MEFs

(B) Bright field representative image of hiPSC colonies on Day 28, after manual selection and transfer to feeder-free culture conditions (scale bar, 200 µm). Clone DEL_4_1.

(C, D, E, F) All hiPSC clones expressed the pluripotency markers NANOG, OCT3/4, TRA-1-60, and TRA-2-49 (scale bars, 100 µm). Clone DEL_5_7.

(G) Extinction of Nanog expression following directed differentiation into endo, meso and ectoderm lineages. Log2 of fold change in RNA abundance (differentiated/undifferentiated)

**Supplementary Figure 2. hiPSCs differentiate into NPCs and neurons**.

(A) Flow cytometric analysis of the transition between Oct4+ pluripotent hiPSCs and early Pax6+ telencephalic neural progenitor cells.

(B) Wild-type (WT), 16p11.2 microdeletion (DEL), and 16p11.2 microduplication (DUP) Day 26 neural rosettes show the typical radially arrayed clusters of neural progenitor cells in brightfield micrographs. Clone IDs for each image: DEL (DEL_5_9, DEL_1_2, DEL_5_8, DEL_9_984), DUP (DUP_3_1, DUP_1_9, DUP_3_3, DUP_1_8), WT (WT_8343.2, WT_8343.2, WT_8343.4, WT_2242.5).

(C) Day 45 immature neurons are characterized by long neurites and expression of the neuronal markers Tuj1 and NeuN (scale bar, 50 µm). Clone WT_8343.5.

**Supplementary Figure 3. Presence of 16p11.2 reprogramming vector integration and transcripts**.

PCR was used to detect reprogramming vector constructs in genomic DNA, and RT-PCR was used to detect mRNA transcripts from each plasmid. Many of the clones were positive for both OCT4 plasmid DNA and transcript. Some clones were also positive for KLF4 or LIN28 plasmid, but no transcripts were detected. A subset of available clones was reprogrammed using additional plasmids carrying EBNA and p53 sequences. No reprogramming vectors were detected in these lines. Plasmid 1 - pCXLE-hOCT4-shp53-F (Addgene plasmid: 27077), plasmid 2 - pCXLE-hUL (L-MyC and Lin28; Addgene plasmid: 27080), plasmid 3 - pCXLE-hSK (Sox2 and KLF4; Addgene plasmid: 27078). Plasmids 4 - pCE-mp53DD (Addgene plasmid: 41856) and plasmid 5 - pCXB-EBNA1 (Addgene plasmid: 41857). Abbreviations: Int, Vector integrant PCR product; Tr, Vector transcription RT-PCR product; ND, not determined; NA, not applicable.

**Supplementary Figure 4. Influence of OCT3/4 integration**.

(A) Total pHH3 counted in Day 26 neural rosette cultures in Int- and Int+ clones normalized by total cell count (p = 0.03174).

(B-E) The first two principal components are plotted and shaded by detection of transcripts against the OCT3/4- bearing plasmid (B), donor genotype (C), donor sex (D), and date of library preparation (E).

(F) Normalized transcript expression levels of neural regionalization candidate genes generated from mRNA-Seq data, as previously described in Figure 3D. hiPSCs are sub-divided into Int+ (red) and Int- (black).

**Supplementary Figure 5. Validation of differentially expressed 16p11.2 interval genes**.

(A) qPCR validation of the 16p11.2 interval genes KCTD13, TAOK2, MAPK3 and SEZ6L2 showing a reduction in gene expression in all the DEL lines. Data are expressed as a log2 fold change compared to the average expression of these genes across all WT hiPSC lines.

(B) Concordance in DE gene fold change among 4 clones that have a microdeletion in 14q11.1, a microduplication in 14q11.2, and a microdeletion in 7p11.2, relative to remaining clones and relative to the average of all DEL clones combined. Each dot represents the value for a given DE gene. DE genes were ranked in order of highest fold increase to largest decrease.

**Supplementary Figure 6. Concordance of differentially expressed gene changes across individual clones**.

Log2(DEL/WT) for normalized counts is plotted for the average across all DEL clones (first panel, red symbols) or individually for each DE gene in each clone. The second panel overlays the average change for all DEL clones (red) on top of all individual values for each DE gene in each DEL clone. Remaining panels plot each DE gene for individual DEL clones. Genes are rank ordered in all plots from largest increase to largest decrease from left to right (the same order is used in all plots). All plots are calculated as DEL values relative to average expression in the combined WT clones.

## ABBREVIATIONS

hiPSC: Human Induced Pluripotent Stem Cells
CNV: Copy Number Variants
SFARI: Simons Foundation Autism Research Initiative
NPC: Neural Progenitor Cells
ASD: Autism Spectrum Disorder
SCZ: Schizophrenia
BD: Bipolar Disorder
ADHD: Attention-Deficit/Hyperactivity Disorder
OCD: Obsessive-Compulsive Disorder
PPD: Phonological Processing Disorder
RELI: Receptive-Expressive Language Impairment
DCD: Developmental Coordination Disorder
CHD: Congenital Heart Disease
GERD: Gastroesophageal Reflux Disease
GAD: Generalized Anxiety Disorder
MDD: Major Depressive Disorder
BMI: Body Mass Index
yrs: Years
MEF: Mouse Embryonic Fibroblasts
SNP: Single Nucleotide Polymorphisms
ICC: Immunocytochemistry
PCR: Polymerase Chain Reaction
WT: Wild-Type
DEL: Microdeletion of the 16p11.2 Locus
DUP: Microduplication of the 16p11.2 Locus
Pax6: Paired Box 6
NCad: N-Cadherin
ZO-1: Zonula Occludens-1
aPKCζ: PAR Complex Protein Atypical Protein Kinase C Zeta
pHH3: Phospho-Histone H3
Tuj1: Class III ß-Tubulin
NeuN: Neuronal Nuclear Protein
SVA: Surrogate Variable Analysis
PCA: Principal Components Analysis
Int+: Integrant-Containing
Int-: Integrant-Free
HUGO: Human Genome Organization
GO: Gene Ontology
NEAA: Non-Essential Amino Acids
hESC: Human Embryonic Stem Cells
DE: Differentially Expressed
FDR: False Detection Rate
miRNA: MicroRNA
GSEA: Gene Set Enrichment Analysis
NIMH: National Institute of Mental Health
SF: Size Factors
VST: Variance Stabilizing Transformation
NA: Not Applicable
ND: Not Determined
WPRE: Woodchuck Hepatitis Virus Posttranscriptional Regulatory Element

## Notes

### Competing Interest Statement

The authors have declared no competing interest.

